# Immuno-informatics approach for multi-epitope vaccine designing against SARS-CoV-2

**DOI:** 10.1101/2020.07.23.218529

**Authors:** Souvik Banerjee, Kaustav Majumder, Gerardo Jose Gutierrez, Debkishore Gupta, Bharti Mittal

## Abstract

The novel Corona Virus Disease 2019 (COVID-19) pandemic has set the fatality rates ablaze across the world. So, to combat this disease, we have designed a multi-epitope vaccine from various proteins of Severe Acute Respiratory Syndrome Corona virus 2 (SARS-CoV-2) with an immuno-informatics approach, validated in silico to be stable, non-allergic and antigenic. Cytotoxic T-cell, helper T-cell, and B-cell epitopes were computationally predicted from six conserved protein sequences among four viral strains isolated across the world. The T-cell epitopes, overlapping with the B-cell epitopes, were included in the vaccine construct to assure the humoral and cell-mediated immune response. The beta-subunit of cholera toxin was added as an adjuvant at the N-terminal of the construct to increase immunogenicity. Interferon-gamma inducing epitopes were even predicted in the vaccine. Molecular docking and binding energetics studies revealed strong interactions of the vaccine with immune-stimulatory toll-like receptors (TLR) −2, 3, 4. Molecular dynamics simulation of the vaccine ensured in vivo stability in the biological system. The immune simulation of vaccine evinced elevated immune response. The efficient translation of the vaccine in an expression vector was assured utilizing in silico cloning approach. Certainly, such a vaccine construct could reliably be effective against COVID-19.

## Introduction

The entire world has been jeopardized due to the COVID-19 pandemic. The accelerated mortality rates have suffused at its extreme in due course of high infectivity rate of this disease. The situation report-180 published on 18th July 2020 (which is the latest till the date of this writing) by The World Health Organization (WHO) has recorded 13,876,441 total infected cases and 593,087 total deaths all over the world^1^. The novel SARS-CoV-2 (as termed by the International Committee on Taxonomy of Viruses) was first reported to cause novel pneumonia in the Wuhan seafood market, China and henceforth an outbreak was observed in the city with a huge number of deaths. The sudden flare-up of the disease due to human-to-human transmission capability of the virus led to a pandemic. Phylogenetic analysis indicated its relation to SARS-like bat viruses which could signify bat as one of the primary reservoirs. This virus has a single-stranded RNA genome ranging from 26 to 32 kilobases in length. It belongs to the β group of coronaviruses. It spreads via respiratory droplets so close contact with infected ones causes acute respiratory distress followed by pulmonary failure in the host. The lineage dates back to Severe Acute Respiratory Syndrome Corona Virus (SARS CoV) of 2003 and Middle East Respiratory Syndrome Corona virus (MERS CoV) of 2012 which exhibited similar characteristics of respiratory distress and alveolar injury^2^. It has been also deciphered that the spike glycoprotein, a structural protein expressed in SARS-CoV-2, has binding affinity to the human Angiotensin Converting Enzyme 2 (ACE2) receptor cells and is responsible for attachment followed by entry into host cells^3^. There are other proteins encoded by viral genome which assist in viral replication and pathogenesis. The current scenario displays emergence of strains from various parts of the world helping out the virus to reach out to more human populations. Although it retards the development of a potential vaccine candidate, the solution to it could be a multi-epitope peptide vaccine, a possible way to combat COVID-19.

The approach of designing this vaccine was entirely in silico. B-cell and T-cell epitopes were predicted for the proteins: membrane glycoprotein, nucleocapsid phosphoprotein, envelope protein, Open Reading Frame (ORF) 6, ORF 7a and ORF 10. The selection of such proteins was due to the following reasons: i) these were found to be 100% conserved among isolated viral strains from India, Italy, USA and China (found with multiple sequence alignment tool) ii) these play an important role in replication, assembly and infection of virus particles. iii) The spike protein was not selected as it did not show 100% conserved sequences among the strains. The nucleocapsid phosphoprotein, membrane glycoprotein and envelope protein are structural proteins of the virus while ORF-6, ORF-7a and ORF-10 are the accessory proteins. The nucleocapsid protein is involved in the stability of viral RNA and processes of the viral replication. The envelope protein is a structural protein playing a significant role in maturation of the virus. The membrane glycoprotein forms a part of the outer viral coat and helps in determining the shape of the viral envelope. This protein can bind to all other structural proteins^4,5^. The ORF 6 protein acts as a signal blocker. It provides blockage to the signals, to the immune system, sent from an infected cell. The ORF 7a helps in release of more viral particles from infected cell. The function remains unexplored for ORF 10 ^6^.

Toxicity as well as resistance to inhibition associated with drug-based therapy is often a hurdle to overcome so peptide vaccines designed with immuno-informatics approach are more optimized and could be developed within a short period. It also paves the way for cost-effective vaccine preparations. Conventional vaccines might consist of several antigenic epitopes of which only a few are required to trigger immune response. The peptide vaccine on the other hand consists of the desired T cell as well as B cell induced immune response. Peptide vaccines are even advantageous than conventional vaccines in providing highly targeted immune response and being devoid of allergenicity^7,8^. The most important aspect to be addressed with a multi-protein vaccine developed with immunodominant proteins from various strains includes broad-spectrum immune response initiated against several strains. The final vaccine construct includes cytotoxic T Lymphocyte (CTL), helper T Lymphocyte (HTL) epitopes, found overlapping with B-cell epitopes, and an adjuvant (β subunit of cholera toxin) at the N-terminal. The adjuvant addition aids as an immune response booster. These epitopes confirm the initiation of both humoral and cell-mediated immune response in the host.

The entire synthesis of the vaccine commenced from the selection of viral proteins on the basis of their conserved sequences among different viral strains. This was achieved with multiple sequence alignment.

This followed the prediction of B-cell and T-cell epitopes for those proteins. The multi-epitope construct was prepared with combination of T-cell epitopes (CTL and HTL) overlapping with B-cell epitopes. The construct with epitopes joined with linkers was found to be antigenic and non-allergenic. It was subjected to prediction of interferon-γ inducing epitopes in it. Then the secondary structure prediction and tertiary structure prediction. The tertiary structure was subjected to refinement along with validation. The vaccine interaction with target immune cell receptors is necessary and the toll-like receptors (TLR) play significant role in activation of innate immunity after recognition of pathogen associated molecular patterns (PAMPs) of viruses and other invading pathogens. Moreover, the TLRs play active role in triggering innate immunity and synchronize adaptive immune response. TLR-2 and TLR-4 are involved in recognition of viral structural proteins followed by production of cytokines and even in SARS CoV, they have been found to induce effective immune responses. Certain studies on SARS CoV and MERS CoV even indicated the noteworthy role of TLR 3 in generating protective responses against the virus^9-14^. So, the vaccine was docked with the immunological receptors: TLR-2, TLR-3 and TLR-4. Molecular dynamics simulations of the vaccine were performed to evaluate in vivo stability of the vaccine. Finally, the codon optimization through in silico approach was performed to ensure its maximal production in the chosen host expression system. The inevitable immune simulation to evaluate immunological response after vaccine introduction in the host. These advanced aspects of computational tools would surely revolutionize vaccine therapeutic studies.

## Results

### Selection of viral protein sequences for vaccine construct

Six viral protein sequences were found to be conserved among all the proteins of SARS-CoV-2 strains isolated from India, USA, Italy and Wuhan (Supplementary Fig. S1). Multiple sequence alignments performed for all the proteins depicted such results. All the viral protein sequences were acquired from NCBI database (https://www.ncbi.nlm.nih.gov/genbank/). The proteins, selected for vaccine construct, comprised of three structural proteins – nucleocapsid phosphoprotein, membrane glycoprotein, envelope protein and three accessory proteins – ORF6, ORF7a and ORF10. These proteins play significant role in varied mechanisms ranging from host cell replication, maturation, assembly of virus particles to viral replication and release of newly formed virus particles. The above-mentioned protein sequences were subjected to further analysis.

### Prediction of B-cell epitopes

Linear B-cell epitopes from all the proteins were predicted using BepiPred 2.0, Bcepred, ABCPRED and SVMTriP web servers. The epitope length was set to 16 amino acids in ABCPRED and 20 amino acids in SVMTriP server. A total number of 148 epitopes from all the proteins were predicted by the servers. 19 of 148 epitopes, were found to be overlapping with the T-cell epitopes which had good prediction scores (Supplementary table S1). Among these 19 epitopes, 7 of them were from nucleocapsid phosphoprotein, 4 were from membrane glycoprotein, 1 each from envelope protein and ORF10, 3 each from ORF6 and ORF7a (Table 1).

**Table 1.**
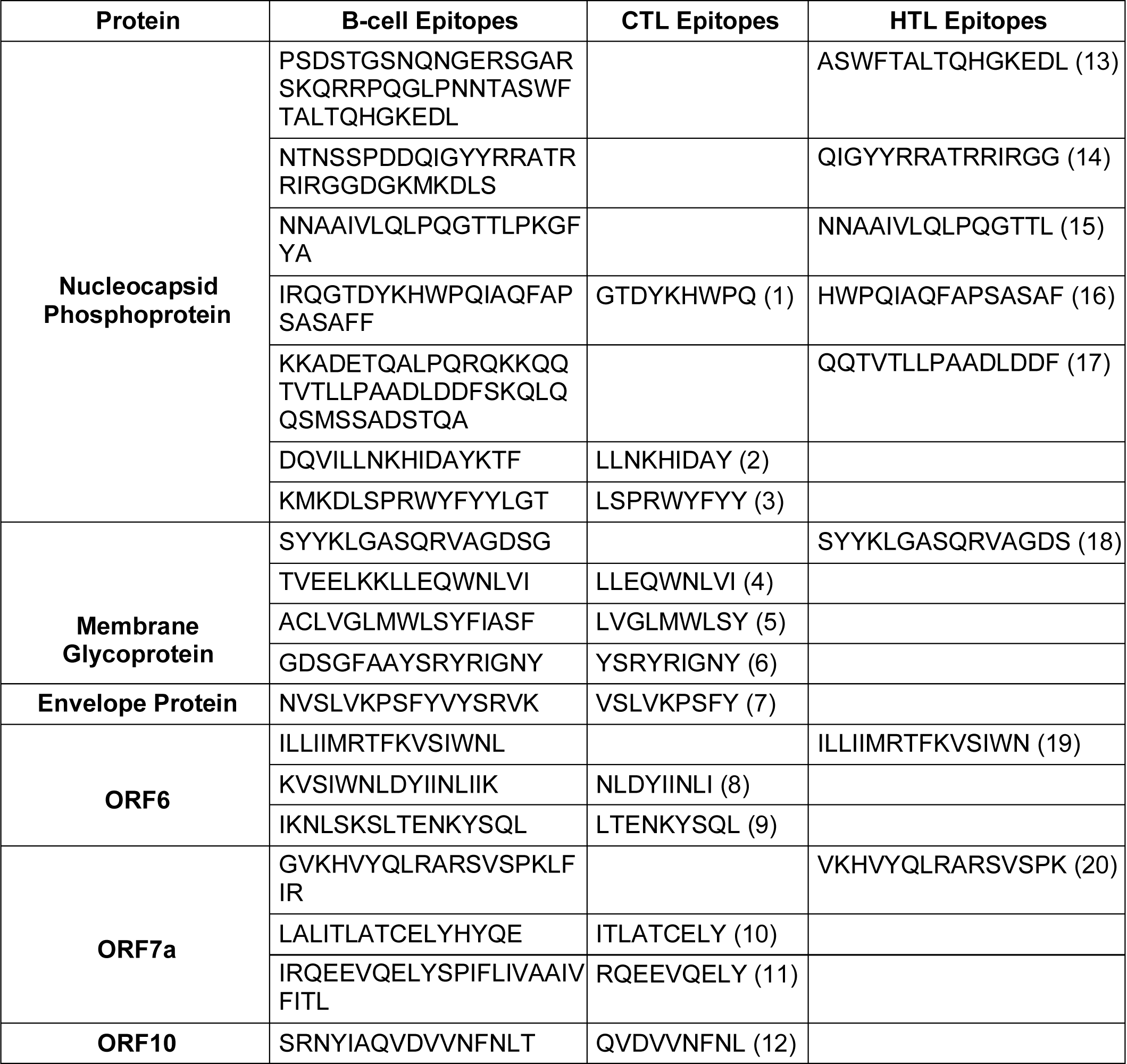
Selected CTL and HTL epitopes overlapping with B-cell epitopes. The numbers in parentheses adjoining epitopes indicates the sequence of epitopes in the vaccine construct from N terminal to C terminal.

### Cytotoxic T Lymphocyte (CTL) prediction

CTL epitopes of A1 supertype were predicted with the NetCTL 1.2 tool with input sequence and 0.15 default weight on C terminal cleavage, 0.05 default weight on TAP transport efficiency along with 0.75 default threshold for epitope identification. 29 epitopes of all the epitopes predicted from the proteins by the server, were marked with ‘<-E’ identifier in output results. The ‘<-E’ marked were assigned as ‘Identified MHC ligands’ by server and they even showed high prediction scores. So, these epitopes were listed down. Out of 29 epitopes, 12 were found to be overlapping with B-cell epitopes and these were finally selected for vaccine construct. Among 12 epitopes, 3 epitopes were from nucleocapsid phosphoprotein, 3 from membrane glycoprotein, 1 each from envelope protein and ORF10, 2 from ORF6 and 2 from ORF7 (Supplementary table S1).

### Helper T Lymphocyte (HTL) prediction

15-mer HTL epitopes having loci in Human Leukocyte Antigen (HLA)-DR, DP, DQ were predicted with the NetMHCII 2.3 tool. The input protein sequences were submitted with default peptide length of 15 amino acids along with the default threshold % rank for strong binder (SB) and weak binder (WB) set to 2 and 10 respectively in the server. Among all the epitopes predicted by the server from all proteins for HLA-DR, 73 ‘SB’ assigned high affinity epitopes were selected as probable epitopes. The epitopes predicted by the server for HLA-DQ and HLA-DR were same in sequence and number, so consideration of epitopes for any one of them was adequate. 8 of these 73 epitopes were found to be overlapping with B-cell epitopes, hence these were selected for vaccine construct. Among these 8 epitopes, 1 each from ORF7a, ORF6, membrane glycoprotein and 5 from nucleocapsid phosphoprotein (Supplementary table S1).

### Multi-epitope vaccine sequence construction

A linear vaccine construct should be capable of satisfying the following criteria i.e. it should possess overlapping HTL and CTL epitopes, it must be immunogenic, antigenic and a non-allergen. Firstly, the overlapping CTL and HTL epitopes whose sequences in turn were found to be overlapping with the B-cell epitopes were selected for final vaccine construct. A total of 12 CTL epitopes and 8 HTL epitopes were selected. The CTL ones were merged by AAY and HTL ones with GPGPG linkers. The β subunit of cholera toxin of 124 amino acids (MIKLKFGVFFTVLLSSAYAHGTPQNITDLCAEYHNTQIHTLNDKIFSYTESLAGKREMAIITFKNGATFQVEVPGSQHIDSQKKAIERMKDTLRIAYLTEAKVEKLCVWNNKTPHAIAAISMAN) was added as an adjuvant at N-terminal of the vaccine construct with EAAAK linker to boost the immune response. After addition of linkers and adjuvant, the final vaccine construct was found to be of length 436 amino acids (Fig. 1A).

**Figure 1.**
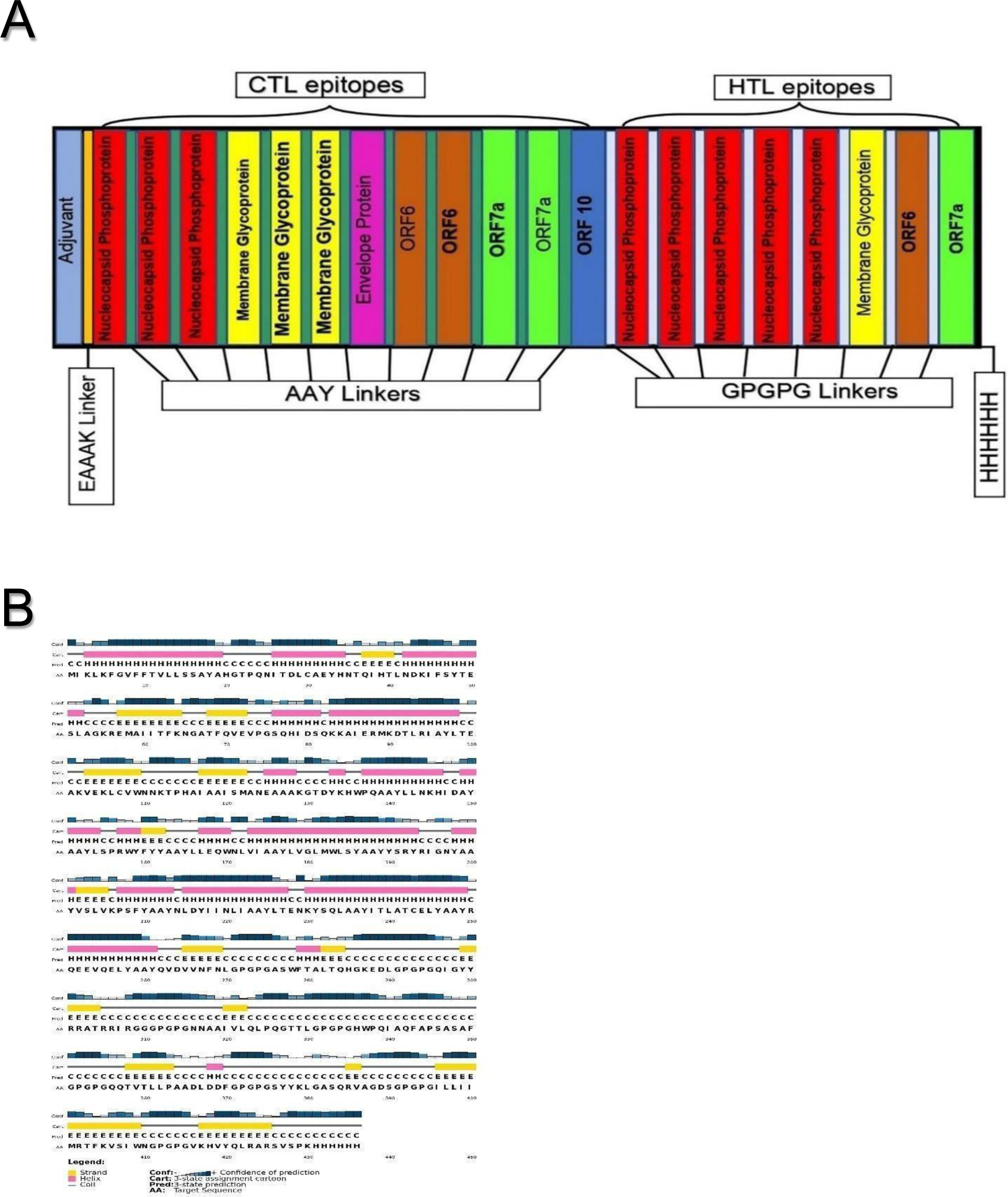
(**A**) Schematic representation of the final multi-epitope vaccine construct. A 436 amino acid long construct with adjuvant at the N-terminal linked with EAAAK linker and 6x-Histidine tag added at the C-terminal CTL epitopes linked with AAY linkers and HTL epitopes linked with GPGPG linkers. (**B**) Schematic representation of the predicted secondary structure of the multi-epitope vaccine construct. The PSIPRED secondary structure prediction results depict the respective contents of α-helix (39%), β-strands (19%) and coils (41%).

### Interferon-γ (IFN-γ) inducing epitope prediction

The IFN-γ inducing epitopes were predicted from the IFN epitope server in the vaccine construct. A total of 428 potential epitopes were predicted by this tool among which 152 showed positive results. This server executes such prediction with the usage of MERCI software.

### Allergenicity and antigenicity prediction of the vaccine candidate

The antigenicity of the final vaccine construct predicted by the VaxiJen 2.0 server was found to be 0.5448 with a virus model at a threshold of 0.4 and with ANTIGENpro it was found to be 0.777. VaxiJen score greater than 0.4 indicates it to be a probable antigen so it depicted its good antigenicity Even the predicted antigen probability score (0.777) in ANTIGENpro adds a support to this fact. The vaccine candidate was predicted to be non-allergenic on both the AllergenFP as well as AllerTOP v. 2.0 servers.

### Physicochemical analysis of vaccine construct

The ProtParam server was used to calculate the physicochemical properties of the vaccine construct. The molecular weight of construct was calculated as 48.14 kiloDaltons (kDa). The theoretical pI (Isoelectric point) was found to be 9.11 which indicates it to be slightly basic in nature. The instability index was found to be 37.77 which confirms it as a stable protein since a score greater than 40 denotes unstable protein. The total no. of positively charged residues and negatively charged residues present in vaccine were found to be 37 and 28 respectively. The estimated half-life of the construct was found to be 30 hours, greater than 20 hours and 10 hours in mammalian reticulocytes (in vitro), yeast (in vivo) and Escherichia coli (in vivo) respectively. The aliphatic index was found to be 84.50 and Grand average of hydropathicity (GRAVY) was found to be −0.172. The aliphatic index score ensures its thermo-stability while the negative GRAVY score confirms its hydrophilicity and it even indicates its ability to interact with solvent molecules. The Recombinant protein solubility prediction tool predicted 100% solubility when overexpressed in *Escherichia coli*.

### Secondary structure prediction

The secondary structure of the multiepitope chimeric peptide vaccine was predicted with PSIPRED tool. It involved 172, 85 and179 amino acids in formation of alpha-helix, beta strand and coil respectively. So, the peptide consists 39% alpha helix, 19% beta strand and remaining 41% as coil (Fig. 1B) (Supplementary Fig. S2). The RaptorX Property server was even utilized to explore solvent accessibility and disordered domain residues in the protein. 42%, 27%, 30% of the residues were found to be exposed, medium exposed and buried respectively. 8% of the residues were found to be in disordered domains.

### Tertiary structure modelling

The tertiary structure of vaccine was modelled with the I-TASSER server. It predicted five tertiary structure models of the chimeric protein based on 10 threading templates of which 1ltrA, 4kxfK, 3chbD, 4l6t and 6b0n were the top five Protein Data Bank (PDB) hits. All the 10 templates (PDB hits) showed good alignment as per their Z-score values (ranging from 1.82 to 5.40). The five predicted models had C-score values in the range −4.08 to −0.64. Since higher score indicates higher confidence the model with higher C-score was selected for further model refinements. Moreover, this model had an estimated TM-score of 0.63±0.13 which is certainly greater than 0.5 and indicates a model of correct topology. These assigned threshold values are independent of protein length. The protein model had an estimated Root Mean Square Deviation (RMSD) value of 8.4±4.5Å. The RMSD value is often sensitive to local error so TM-score has been estimated (Fig. 2A).

**Figure 2.**
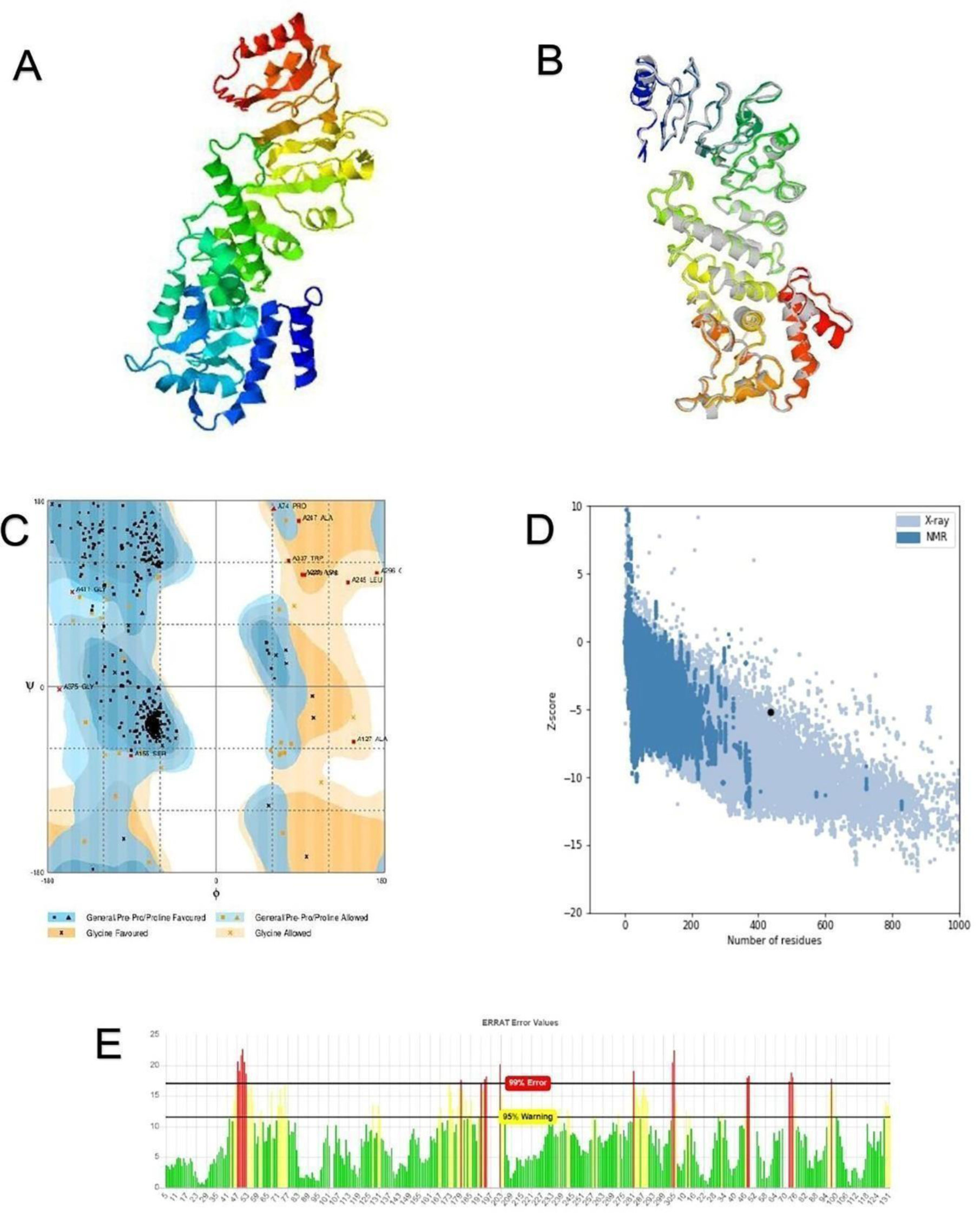
Modelling, Refinement and validation of the multi-epitope vaccine construct model. **(A)** The predicted tertiary structure of the vaccine construct. **(B)** Tertiary structure of the vaccine after model refinement. Refined regions in the model are indicated with grey colour. **(C)** Validation of vaccine tertiary structure by Ramachandran plot analysis indicating 90.3% residues in favored region, 7.1% residues in allowed region and 2.5% residues in outlier region. **(D)** Validation of model with ProSA-web providing a z score of −5.14. The black coloured spot in the plot indicates z score. **(E)** Validation by ERRAT with quality factor 80.73.

### Tertiary structure refinement

The predicted tertiary (3D) structure of vaccine construct was further subjected to model refinement using GalaxyRefine server. GalaxyRefine generated 5 models after refinement, out of which model 1 (Fig.2B) was selected for having best scores compared to others depending on various parameters comprising of GDT-HA (0.948), RMSD (0.433), MolProbity (2.492), Clash score (19.8), Poor rotamers (1.5) and 88.9% favoured region in Ramachandran plot.

### Tertiary structure validation

The validations of the predicted tertiary structures were performed with RAMPAGE server performing Ramachandran plot analysis of the modelled vaccine construct. The results showed 90.3%, 7.1% and 2.5% of the residues of the protein in favored, allowed and outlier regions respectively (Fig. 2C). The favoured region score is almost consistent with the score of refined models selected in GalaxyRefine. The determination of quality and potential errors was inevitable so the ProSA-web and ERRAT servers were utilized respectively. The chosen refined model showed up an overall quality factor of 80.73% in ERRAT server (Fig. 2E) while ProSA-web gave a Z-score of −5.14 for the input vaccine model. The obtained Z score lies within the range of commonly found native proteins of comparable size so the purpose of validation gets fulfilled with the acquired results (Fig. 2D).

### Prediction of discontinuous B-cell epitope

ElliPro server which predicts epitopes on the basis of tertiary structures was the medium of prediction of Discontinuous B-cell epitopes. Ten epitopes were predicted from the server of which the epitope (of 75 residues) with maximum score 0.755 was predicted as a discontinuous epitope (Supplementary Table S2). 14 linear B-cell epitopes were even predicted by the server (Supplementary Table S3).

### Molecular docking of final vaccine construct with immunological receptors

ClusPro online protein-protein docking server was used to perform molecular docking between refined vaccine model and immune receptors TLR-2 (PDB ID-6NIG), TLR 3 (PDB ID-2A0Z) and TLR 4 (PDB ID-4G8A). The output results displayed 30 clusters for each docked complex ranked (0-29) according to cluster members. The weighted scores of energies of the clusters were also provided. Cluster 1 of TLR-2 – vaccine docked complex (Supplementary Fig. S3) with 32 members having lowest energy of −1197.2 and Cluster 0 of TLR-3 – vaccine docked complex (Supplementary Fig. S3) with 97 members (highest no. of members) having lowest energy of −1524.4, were selected for further analysis. Cluster 22 of TLR-4 docked complex showed lowest energy weighted score but it had only 12 members so cluster 2 (Supplementary Fig. S3) with 48 members, having lowest energy of −1207.7 was selected. The interaction surface residues of docked complexes were visualized with BIOVIA Discovery Studio Visualizer software (Fig. 3)

**Figure 3.**
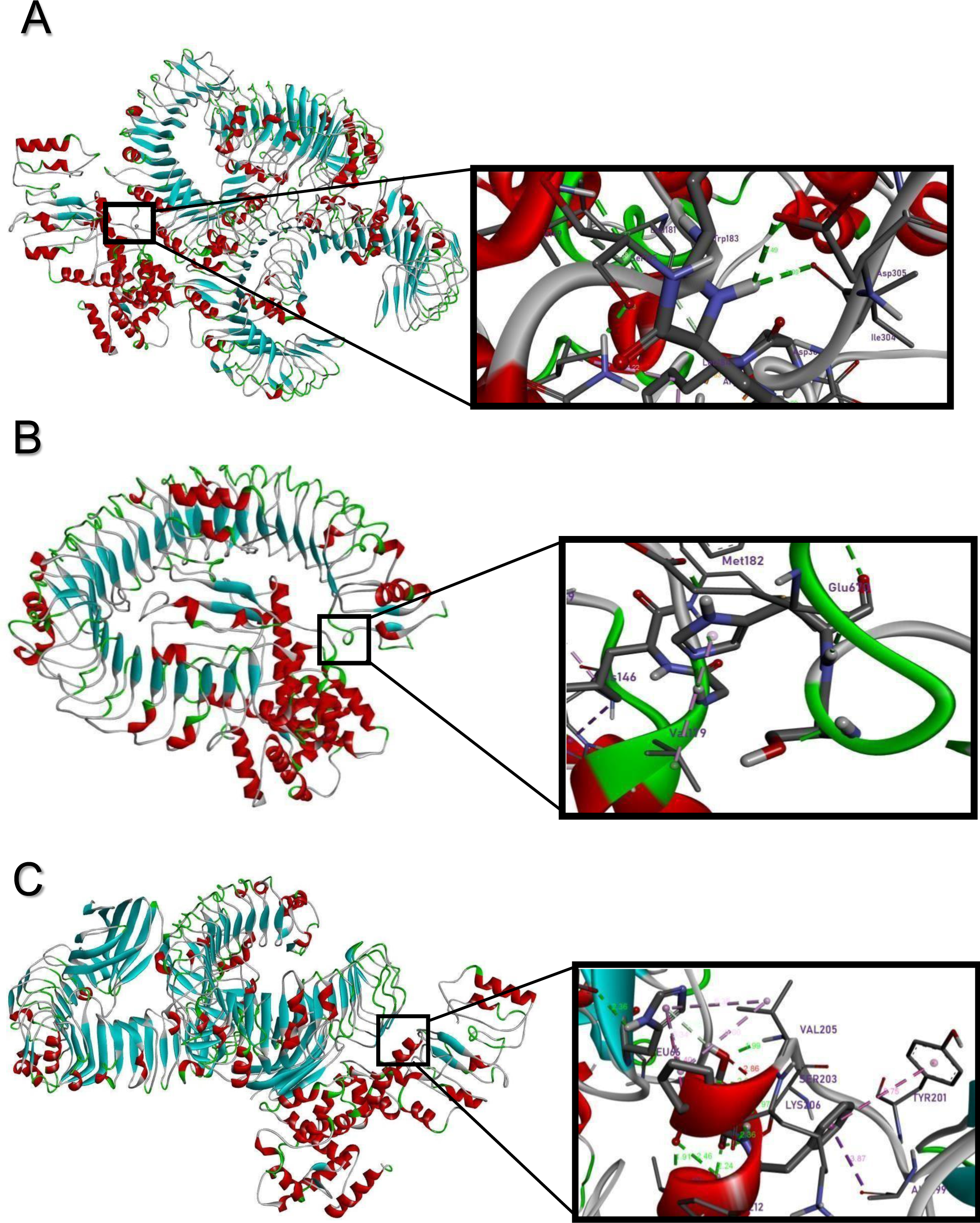
Visualization of interacting residues of docked complex. (**A**) TLR2-vaccine complex (**B**) TLR3-vaccine complex (**C**) TLR4-vaccine complex. The beta-strands, alpha-helix, loops, coils are indicated in the proteins of docked complex with colours blue, red, green and grey respectively.

### Analysis of binding affinity of docked complexes

The study of binding energetics is inevitable for biological complexes. The binding affinity of a complex or the Gibbs free energy (ΔG) i.e. binding free energy determines the probability of occurrence of interactions at specific cellular conditions. So, the PRODIGY web server was used to analyse the binding affinity of the 3 docked complexes. The input accepts PDB file of the docked complexes along with the interactor chains of receptor and ligand. The temperature was set to default value of 25°C. The ΔG values obtained for the vaccine-TLR2, vaccine-TLR3, vaccine-TLR4 complex were found to be − 18.9 kcal mol^−1^, − 19.1 kcal mol^−1^ and − 18.0 kcal mol^−1^ respectively. The results evinced energetically feasible docking, as depicted by the negative values of Gibbs free energy. The dissociation constants of the complexes, number of interfacial contacts per property and non-interacting surface per property are shown in Table 2.

**Table 2.** Results for binding energetics. Predicted results of binding affinity (Gibbs free energy), dissociation constant, number of interfacial contacts per property, non-interacting surface per property by PRODIGY web server.

| Docked Complexes | Gibb's Free Energy (kcal/mol) | K<sub>d</sub> (M) at 25° C | Number of Interfacial Contacts (ICs) per property |  |  |  |  |  | Non- Interacting Surface (NIS) per property |  |
| --- | --- | --- | --- | --- | --- | --- | --- | --- | --- | --- |
|  |  |  | ICs charged - charged | ICs charged-polar | ICs charged-apolar | ICs polar-polar | ICs polar-apolar | ICs apolar-apolar | NIS charged (%) | NIS apolar (%) |
| Vaccine-TLR2 | -18.9 | 1.3E-14 | 10 | 18 | 49 | 4 | 33 | 40 | 26.42 | 31.62 |
| Vaccine-TLR3 | -19.1 | 1.0E-14 | 28 | 25 | 67 | 2 | 17 | 19 | 21.29 | 36.13 |
| Vaccine-TLR4 | -18.0 | 6.1E-14 | 7 | 6 | 34 | 9 | 42 | 33 | 23.20 | 35.03 |

### Molecular Dynamics simulation of the final vaccine construct

Molecular dynamics simulation (MDS) results help to identify the stability of a protein under in vivo conditions. MDS was performed using Galaxy server which in turn uses the GROMACS engine for running simulations. Solvated and neutralized vaccine construct was energy minimized before the simulation. As soon as the force reaches a value < 1000 kJ/mol, the protein becomes energy minimized. Average temperature of the system was set at 300.17 K with a drift of 2K after NVT equilibration for 50,000 steps with 0.002 ps time step. Average density of the system calculated was 1057.95 kg/m^3^ with a drift of 5.05 kg/m^3^. Plot of pressure evaluation shows fluctuation in pressure throughout the NPT equilibration phase while maintaining an average value close to 1 bar. Trajectory analysis after production simulation of 10ns was performed to validate the protein stability inside biological system. Root Mean Square Deviation (RMSD) backbone plot having mild fluctuations indicate the stability of the vaccine construct over time. The observed high peaks of Root Mean Square Fluctuation (RMSF) plot of sidechains suggests high degree of flexibility of the vaccine (Fig. 4). Principal Component Analysis (PCA) and Dynamic Cross-Correlation matrix (DCCM) plots are also shown in Fig. 4.

**Figure 4.**
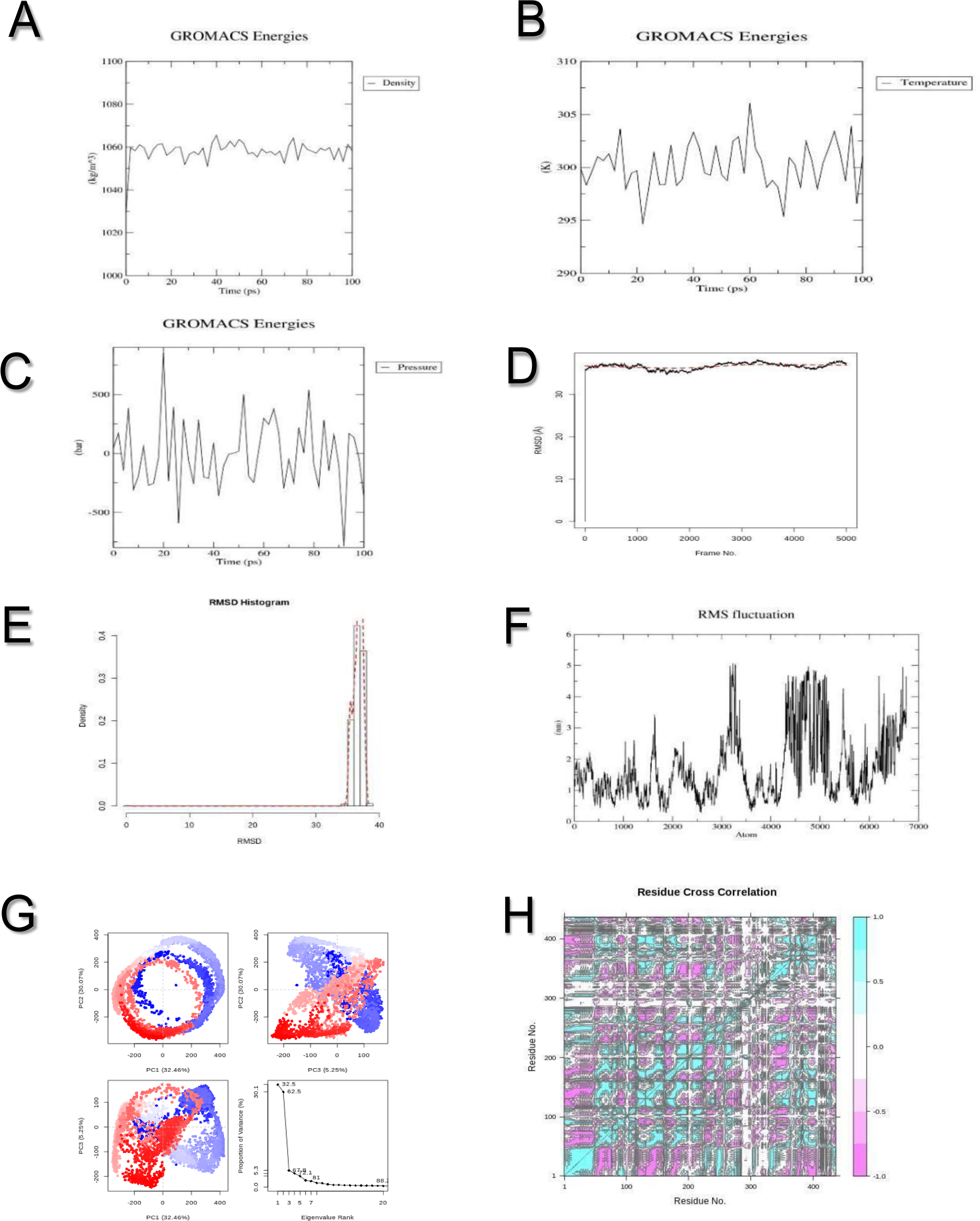
Molecular Dynamics Simulation results of the vaccine construct. (**A**) Density evaluation plot indicating an average value of 1057.95 kg/m^3^ having a drift of 5.05 kg/m^3^ (**B**) After 50,000 steps of NVT equilibration for 100 ps, the average value of temperature is stabilized at 300.17 K with a drift of 2 K (**C**) Pressure plot depicts fluctuations in pressure throughout the NPT equilibration phase with an average value close to 1 bar. (**D**) RMSD plot of backbone showing mild fluctuation (approx. 3.5 nm) indicating in vivo stability of vaccine. (**E**) RMSD histogram validating RMSD plot. (**F**) RMSF plot of sidechains revealing high degree of fluctuation confirming high flexibility of vaccine construct. (**G**) PCA representation for vaccine construct’s MD simulation. Corresponding eigenvalue (PCs) signify the respective content of the total mean square displacement of positional variations of residues as included in each dimension. Periodic transitions among the conformers via trajectory is indicated with the uninterrupted colour scale from blue to white and then to red. (**H**) DCCM for vaccine construct based on C-alpha atoms. Blue colour indicates positive correlation and pink colour indicates negative correlation in respective motion of residues.

### Protein structure validation after Molecular Dynamics Simulation

Ramachandran plot analysis by RAMPAGE shows a significant decrease in the percentage of residues (2.1%) in outlier regions which was 2.5% before the MD simulation. Although, there is a decrease in percentage of residues in favoured regions (83.8%), a significant increase in percentage of residues (14.1%) lying in allowed regions was noted (Supplementary Fig. S4). Before MD simulation, 90.3% and 13% residues were present in favoured and allowed regions respectively. ProSA – web shows a Z-score of −4.58 which lies within the range of native proteins of comparable size (Supplementary Fig. S4). ERRAT quality factor is decreased from 80.73 to 64.74 which is much less than the rejection limit (> 95%). Moreover, a result of >50 indicates a protein of good topology (Supplementary Fig. S4). So, it could be concluded that in spite of certain changes in structure after MD simulation, the vaccine construct proves to be stable in vivo within biological system.

### Codon optimization and in silico cloning of the vaccine construct

In silico cloning of the vaccine construct is of immense noteworthy for its expression in *Escherichia Coli* expression system. As a result, the codon optimization of construct is inexorable as per usage in expression system in order to assure efficient translation. The Java Codon Adaptation Tool (JCat) was utilized for codon optimization of the final vaccine construct for maximal protein expression in *Escherichia Coli* (K-12 strain). The length of the generated cDNA sequence after codon optimization was of 1308 nucleotides. The optimum range for Codon Adaptation Index (CAI) of the optimized nucleotide sequence is greater than 0.8 and for the vaccine, it was found to be 0.96 which indicates high expression of gene. The average GC content of the adapted sequence was 53.98% which also indicates the possibility of good expression of the vaccine candidate in the host system since the optimal percentage of GC content lies in the range of 30-70%. Finally, the design of the recombinant plasmid was accomplished in silico by inserting the adapted codon sequences into pET-28a (+) vector using SnapGene software (Fig. 5). This study ensured effective cloning strategy of the multi-epitope vaccine construct.

**Figure 5.**
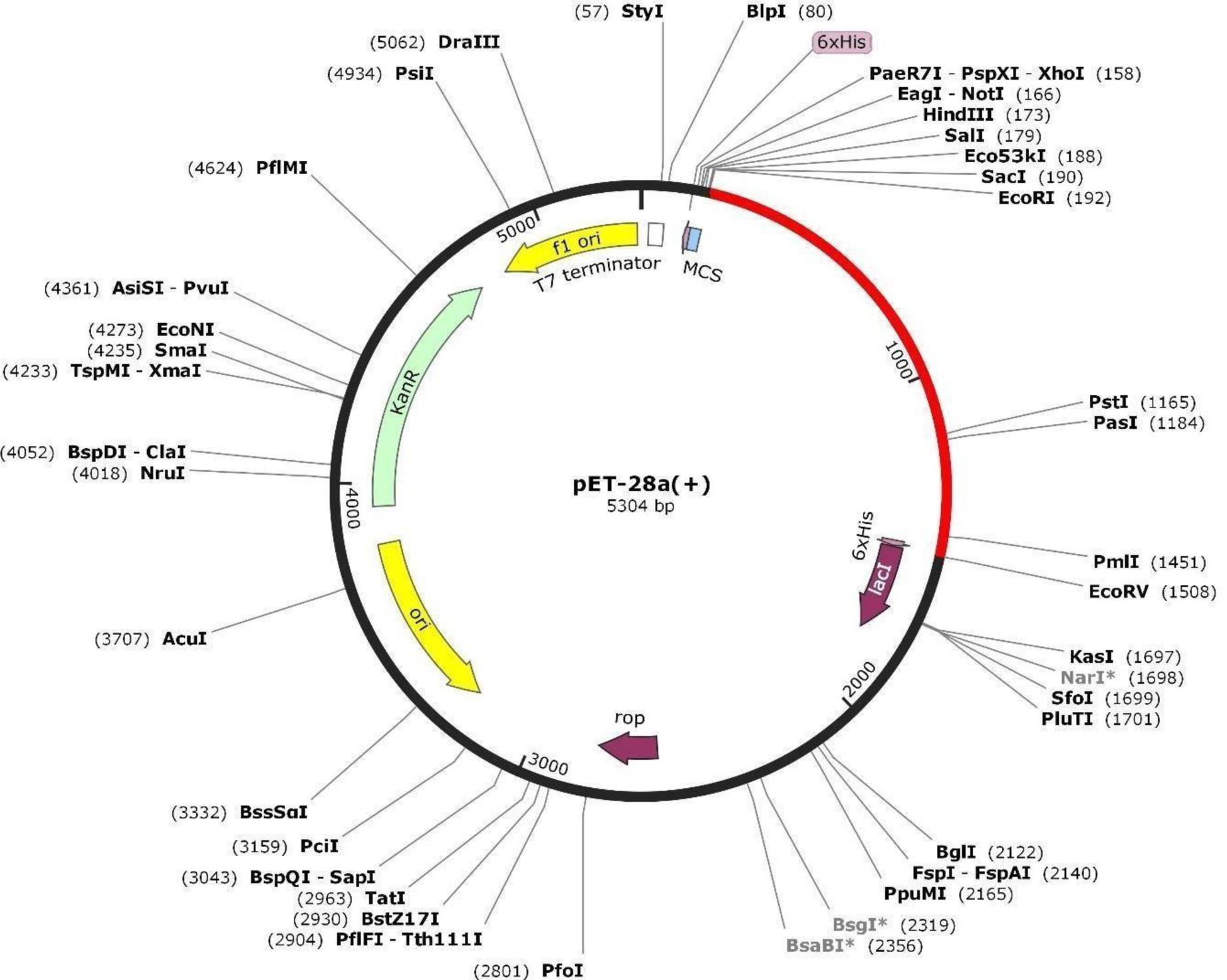
In silico restriction cloning of the final vaccine construct sequence into the pET28a (+) expression vector. The red part represents the gene encoding for the vaccine and the black circle represents the vector backbone.

### Immune Simulation

The immune simulation of the vaccine was performed with C ImmSim server. The results depict secondary and tertiary immune response (IgG1, IgG2, IgG + IgM) to be greater than primary immune response (IgM). The antigen concentration decreases and immunoglobulin concentration (IgM, IgG1+ IgG2, IgG + IgM) increases after vaccine injection. Long lasting B cells exhibit isotype switching ability and development of memory cells. T_H_ and T_C_ cell responses are found high with corresponding development of memory. The pre-activation of T_C_ cell response is found during vaccination. Natural Killer cells and Dendritic Cells show consistent response throughout. High levels of macrophage activity are also indicated. 12 doses of injections consistently given 4 weeks apart show high levels of IFN-gamma and Interleukin (IL)- 2 elicited which show consistency with the prediction of IFN-gamma epitopes in the vaccine. Two aspects of input were implemented for immune simulation. One of the incorporated methods was that after vaccination, the live replicating virus was simulated at day 366. No antigenic surge was present in this case and the antigen was contained immediately. It reveals the presence of protective antibodies. The second aspect was that without prior vaccination, live replicating virus was simulated at around similar day (366). The antigenic surge present in this case indicates the failure to contain the virus in spite of presence of a mild immune response (Fig. 6). (Supplementary Fig. S5)

**Figure 6.**
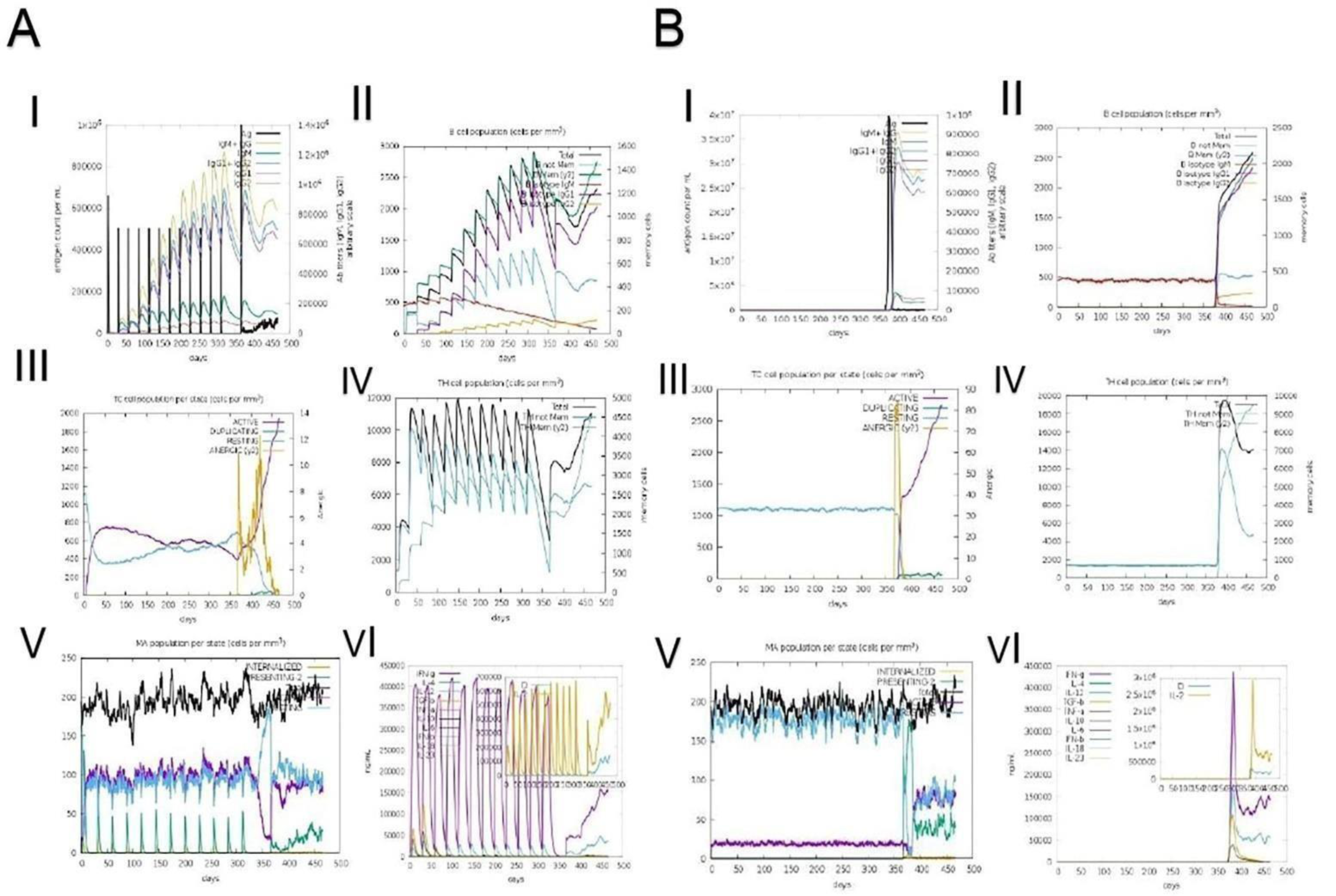
**(A)** 12 doses of vaccine injections were given for almost 15 months and a live replicating virus was injected at around day 366 **(Ai)** indicates the increase of antigen concentration and relative antibodies responses. Live-replicating virus, injected two months after last vaccine dose is contained immediately due to the production of protective immunoglobulins highlighting the efficiency of the vaccination. **(Aii)** represents the count of B-cell population. **(Aiii-iv)** indicating the activation of cytotoxic T-cells and helper T-cells. (**Av**) Macrophage activation is shown **(Avi)** High levels of IFN-gamma, Tumor Necrosis Factor (TNF)-b, Interleukin (IL)-12 and IL-2 indicates good immune response. **(B)** Comparison with a control experiment where the live virus is injected at the same time period (at around day 366) but without prior vaccination in this case. **(Bi)** Antigenic surge for a longer period of time indicates the absence of any memory cells without prior vaccination. **(Bii-vi)** absence of strong immune response due to lack of vaccination could not contain the antigenic load which further proves the efficiency of the vaccine construct.

## Discussion

The development of a multi-epitope vaccine candidate in silico plays a more significant role in therapeutics as compared to single epitope since it facilitates to identify antigenic epitopes which could initiate targeted immune response. Moreover, the immune-informatics approach of developing such candidate secures cost effectiveness and time-saving methods to prepare the vaccine construct^12-13^. We aimed to develop a multi-protein, multi-epitope vaccine construct which could trigger humoral as well as cell-mediated immune response when injected in the host since the construct contains both T_C_-cell and T_H_-cell epitopes, both overlapping with B-cell epitopes. The designed vaccine would surely be able to provide double protection in targeted immunological aspects. The vaccine candidate consists of epitopes for multiple viral proteins which could provide noteworthy defence against infection. The design of this construct commenced with selection of six proteins with conserved sequences among 4 different SARS-CoV-2 strains. The B-cell, CTL and HTL epitopes were predicted for the selected proteins. The CTL and HTL epitopes, which overlapped with B-cell epitopes, were joined by linkers and utilized to construct this vaccine. An adjuvant was even added to the N terminal of this construct to boost the immune response. The 6x-Histidine (His) tag was added at the C-terminal for serving purification purposes. The vaccine was found to be non-allergic and antigenic. Interferon-γ epitopes were even identified in the vaccine which confirmed the potential of vaccine to initiate functioning of Interferon-γ exhibiting immune-stimulatory actions. The secondary structure of the vaccine was predicted and even the tertiary structure was modelled. This 3D model was further refined and validated. The vaccine was docked with TLR2, TLR3 and TLR4 receptors. These receptors are able to trigger innate immune response. The binding affinity related parameters of docked complexes were further analysed. The binding free energy of all vaccine-receptor complexes were found to be highly negative which confirmed most stable binding and the dissociation constants were also found to be of low values for all three complexes. The results obtained after molecular dynamics simulation of the vaccine assured in vivo stability of the construct. The results for validation of tertiary structure of the construct after MD simulation showed the stability in structure. The in-silico cloning approach was performed to ensure maximal expression of the vaccine in an expression vector. The immune simulation was finally performed for the evaluation of immunogenic profile of the vaccine. Antigen is contained immediately after infection with live virus. Antigen concentration is found to be decreasing immediately due to the presence of memory cells since prior vaccination helped in memory cell development. The memory B cells, memory helper T and memory cytotoxic T cells are triggered in presence of the live virus which ultimately contain the virus. On the other hand, without prior vaccination and memory cells being absent, caused antigenic surge for a much longer amount of time after the injection of the live virus.

## Methodology

### Accession to viral protein sequences

All the protein sequences of SARS-CoV 2 strains of India, Italy, USA and Wuhan, were retrieved from NCBI (https://www.ncbi.nlm.nih.gov/genbank/) with Reference Sequence ID MT050493.1, MT066156.1, MN985325.1 and NC_045512.2 respectively^15^.

### Identification of proteins for vaccine development

Six protein sequences were found to be 100% conserved among different strains. It was done using Multiple Sequence Alignment tool: Clustal Omega (https://www.ebi.ac.uk/Tools/msa/clustalo/) which incorporates progressive approach, seeded guide trees and HHalign package for generating swift and accurate alignments for three or more input sequences^16^.

### B-cell epitope prediction

The B-cell epitopes for the six proteins were predicted with four web servers. Firstly, BepiPred2.0 (http://www.cbs.dtu.dk/services/BepiPred/) which relies on random forest algorithm for discerning epitope database annotated from various antigen-antibody complexes^17^. Secondly, Bcepred (https://webs.iiitd.edu.in/raghava/bcepred/bcepred_submission.html) which is solely based on physico-chemical properties of dataset created with epitope sequences acquired from Bcipep and SWISS-PROT database^18^. The default threshold was set for all the parameters, hydrophilicity (2), accessibility (2), exposed surface (2.4), antigenic propensity (1.8), flexibility (1.9), turns (1.9), polarity (2.3) and combined (1.9) as input. Thirdly, ABCPRED(https://webs.iiitd.edu.in/raghava/abcpred/ABC_submission.html) which predicts on the basis of artificial neural network methodology using fixed length patterns^19^. The default threshold 0.51 and the default overlapping filter was set on for input. Fourthly, SVMTriP (http://sysbio.unl.edu/SVMTriP/prediction.php) which implements a new method to predict antigenic epitope with sequences from the Immune Epitope Database (IEDB). This server utilizes Support Vector Machine (SVM) combining the Tri-peptide similarity and Propensity scores (SVMTriP) in order to achieve better prediction performance^20^. The default epitope length was set to 20 amino acids.

### Cytotoxic T lymphocytes (CTL) epitope prediction

The CTL epitopes for the peptide vaccine were acquired from a freely accessible NetCTL 1.2 (http://www.cbs.dtu.dk/services/NetCTL/) server with default threshold value for epitope identification set at 0.75. This tool incorporates prediction of Major Histocompatibility Complex (MHC) class I binding peptides, proteasomal C-terminal cleavage, and TAP (Transporter Associated with Antigen Processing) transport efficiency. These epitopes are recognized by Major HistoCompatibility Complex Class I supertype A1, found in human population. MHC class I binding and proteasomal cleavage are performed using artificial neural networks while a weight matrix is used to determine TAP transport efficiency^21^. The default threshold value for epitope identification set at 0.75 was used for the prediction of CTL epitopes.

### Helper T-cell (HTL) epitope prediction

HTL epitopes of 15-mer length for human alleles were predicted by the Net MHCII 2.3 Server (http://www.cbs.dtu.dk/services/NetMHCII/). The server helps to predict peptides binding to HLA-DR, HLA-DQ, and HLA-DP alleles with the aid of artificial neuron networks. Each epitope is assigned IC50 value to deduce receptor affinity ^22,23^. High-affinity peptides are supposed to possess IC50 values <50 nM. IC50 value lesser than 500 nM indicates intermediate affinity and values less than 5000 nM indicate low affinity. So, on one hand the percentile rank may therefore be inversely related to the epitope binding affinity while on the other hand directly related to the IC50.

### Construction of a multi-epitope vaccine sequence

The HTL and CTL epitopes which overlapped with linear B-cell epitopes, were utilized to prepare the construct. Effective functioning of each epitope is ensured with insertion of GPGPG and AAY linkers used to link all the epitopes of HTL and CTL respectively^24^. The amino acid sequence of the β-subunit of cholera toxin was added as an adjuvant at the N-terminal of the construct with the help of EAAAK linker^24^.

### IFN-γ inducing epitope prediction

Interferon gamma (IFN-γ) provides an amplified response to MHC antigens, stimulates macrophages and natural killer cells playing an active role in adaptive and innate immune response. IFN-γ epitopes were predicted from IFNepitope server(http://crdd.osdd.net/raghava/ifnepitope/predict.php). The server essentially constructs overlapped sequences utilized for epitope prediction. The server compiles all MHC Class II binders from Immune Epitope Database and Analysis resource (IEDB) and categorizes them into inducing and non-inducing binders. The algorithm of this server runs with three models-Motif based model, Support Vector Machine based model and Hybrid approach of both SVM and Motif based, for an input query peptide^25^. The IFN-γ epitopes in construct were predicted with Hybrid approach.

### Antigenicity and allergenicity prediction

The antigenicity was predicted by alignment-free online server ANTIGENpro (http://www.scratch.proteomics.ics.uci.edu/). It remains alignment-free and does not depend on any pathogen identity for antigenicity prediction. A two-step process based on five algorithms is implemented in the prediction. SVM classifier generates a brief result of prediction informing about the probability of a peptide possessing characteristics of an antigen^26^. VaxiJen v. 2.0 server (http://www.ddg-pharmfac.net/vaxijen/VaxiJen/VaxiJen.html) validated antigenicity with the default set to 0.4 and target organism as virus. It engrosses auto cross covariance (ACC) transformation of protein sequences into uniform vectors of principal amino acid properties^27^. The allergenicity of construct was predicted with online tool AllerTOP 2.0 (https://www.ddg-pharmfac.net/AllerTOP/). It incorporates the method based on auto cross covariance (ACC) transformation of protein sequences into uniform equal-length vectors. The principal properties of the amino acids were represented by five E-descriptors: amino acid hydrophobicity, molecular size, helix-forming propensity, relative abundance of amino acids, and β-strand forming propensity. The proteins are categorized using k-nearest neighbor algorithm (kNN, k=1) based on training set which consists of 2427 known allergens and 2427 non-allergens^28^. The allergenicity was further validated with AllergenFP v.1.0 (http://ddg-pharmfac.net/AllergenFP/) working on a four-step algorithm. Firstly, the amino acids in the protein sequences in data sets were described by five E-descriptors: amino acid hydrophobicity, molecular size, helix-forming propensity, relative abundance of amino acids, and β-strand forming propensity. and the generated strings were transformed into uniform vectors by auto-cross covariance (ACC) transformation. The vectors were transformed into binary fingerprints and compared in terms of Tanimoto coefficient^29^.

### Prediction of various physicochemical properties

The physicochemical properties of the vaccine like molecular weight, extinction coefficient, amino-acid composition, estimated half-life, no. of amino acids, aliphatic index, instability index, theoretical pI and grand average of hydropathicity were explored using ProtParam tool (https://web.expasy.org/protparam/) ProtParam implements the N-end rule to predict half-life, assigns weight value of instability to dipeptides for instability index, mole-percent as well as volumes occupied by aliphatic amino acid side chains for aliphatic index and average of hydropathy values for GRAVY score^30^. The Recombinant protein solubility prediction tool (https://biotech.ou.edu/) was used to predict protein solubility considering the assumption that it is over expressed in *Escherichia coli*. A statistical model has been incorporated in this tool which was built using logistic regression of32 possible parameters^31^.

### Secondary structure prediction

The secondary structure predictions of vaccine was performed with PSIPRED server(http://bioinf.cs.ucl.ac.uk/psipred/). The accurate prediction is accomplished with incorporation of two feed-forward neural networks performing synchronized analysis of output obtained from first network followed by output analysis of the input from initial network prediction generated from PSI-BLAST (Position Specific Iterated – BLAST)^32^. The RaptorX Property Prediction tool (http://raptorx.uchicago.edu/StructurePropertyPred/predict/) was utilized to analyse solvent accessibility of the vaccine construct. The solvent accessibility (ACC) and disorder regions (DISO) are predicted with a strong method of in-house deep learning model, Deep Convolutional Neural Fields (DeepCNF) incorporated in the server. On one hand, the DeepCNF is involved in modelling the complex sequence–structure relationship by a deep hierarchical architecture and on the other hand models interdependency between adjacent property labels. The following characteristics of around 66% Q3 accuracy for 3-state solvent accessibility are attained by the server and approx. 0.89 area under the ROC curve (AUC) for disorder prediction is even accomplished by the server ^33-36^.

### Tertiary Structure prediction

The tertiary structure prediction of construct was performed with Iterative Threading Assembly Refinement (I-TASSER) on-line server (https://zhanglab.ccmb.med.umich.edu/I-TASSER/) which utilizes hierarchal approach to cast light on protein structure. It accepts input protein sequence and proceeds with identification of structural templates from the PDB by multiple threading approach Local Meta-Threading Server (LOMETS), along with iterative template-based fragment assembly simulations generated full-length atomic models. Function predictions of the target are then executed by re-threading the 3D models by means of protein function database BioLip^37^. The output results display five predicted tertiary structure models of the input sequence based on the first 10 PDB hits as the threading templates. The five models are also provided with C-scores and TM-scores.

### Tertiary Structure Refinement

The refinement of the template-based protein tertiary structure is unavoidable for achieving precision in model. The refinement was performed with the freely accessible GalaxyRefine server (http://galaxy.seoklab.org/cgi-bin/submit.cgi?type=REFINE). The refinement method improves the global and local structural qualities. This server operates firstly via rebuilding all side-chains by placing the highest-probability rotamers. The next rotamers of highest probabilities are attached on having encounter with steric clashes. The new model with rebuilt sidechains is subjected to refinement with mild and aggressive relaxation methods^38^. The model 1 with lowest energy is generated by mild relaxation and the other four models are generated by aggressive relaxation. The triaxial loop closure method is applied in order to avoid breaks in model structures caused by perturbations to internal torsion angles. The GalaxyRefine displays an output of five generated structure models and their respective GDT-HA, MolProbity, Clash, RMSD, Poor rotamers’ scores alongwith % favoured regions in Ramachandran plot.

### Tertiary Structure Validation

The tertiary structure validations of refined models were performed with RAMPAGE, ERRAT and ProSA-web servers. The RAMPAGE (http://mordred.bioc.cam.ac.uk/~rapper/rampage.php) server accepts PDB input of the model and displays favoured, allowed and outlier region residues in Ramachandran plot along with diagrammatic representation^39^. This server generates the plot with energetically favourable dihedral angles of the amino acids which are calculated on the basis of the Van der Waal radius of the sidechain. The ERRAT (https://servicesn.mbi.ucla.edu/ERRAT/) performing statistical analysis of the non-bonded atomic interactions, involves a quadratic error function for characterization of pair-wise atomic interactions from nine-residue sliding windows accumulated in a database of relied protein structures. The error prone regions are then recognized with pattern of interactions from each window^40^. The ProSA-web server (https://prosa.services.came.sbg.ac.at/prosa.php) involves a computational engine which incorporates knowledge-based potentials of mean force for evaluation of model accuracy. The output displays model *z*-score and a plot of residue energies. The generated *z*-score indicates overall model quality and measures its deviation from the total energy of the structure with respect to an energy distribution derived from random conformations. All calculations are carried out with C^α^ potentials which enables its applications to low-resolution structures^41,42^.

### Prediction of discontinuous B-cell epitopes

Discontinuous B-cell epitopes of the vaccine candidate were predicted with ElliPro web server (http://tools.iedb.org/ellipro/). The server takes PDB file input and tries to find the protein or its homologues in PDB with the protein BLAST. It executes three algorithms for performing the following tasks; firstly, protein shape is approximated as an ellipsoid, secondly, the residue protrusion index is calculated and thirdly, the neighbouring residues are clustered based on their Protrusion Index values. The output results generate a score for each epitope predicted which is based on Protrusion Index value averaged over epitope residues. The results even display scores, provide access to 3D structure visualization of predicted linear epitopes as well as the discontinuous ones followed by 2D score chart of residues in input. This method involves detection of protein’s 3D shape by means of quantitative approximation of no. of ellipsoids. The epitope prediction parameters, minimum score and maximum distance are set to default score 0.5 and 6 respectively^43^.

### Molecular Docking of vaccine construct with immunological receptor

The immunoreceptor-vaccine interaction is highly significant to elicit the immune response. So, it was inexorable to perform molecular docking of the vaccine construct with immunological receptor molecules, TLR-2, TLR-3 and TLR-4. The docking was executed with ClusPro 2.0 protein-protein docking online server (https://cluspro.bu.edu/login.php). It accepts PDB file input of the two proteins to be docked. The server incorporates the following three computational steps, firstly it performs rigid body docking by means of sampling billions of conformations, secondly root-mean-square deviation (RMSD) based clustering of the 1000 lowest energy structures generated in quest of the largest clusters which are expected to represent the most likely models of the complex and thirdly, it executes refinement of selected structures with the help of energy minimization. The rigid body docking runs with PIPER docking program. This program utilizes Fast Fourier Transform correlation approach. PIPER indicates the interaction energy between two proteins. The second step involves clustering of lowest energy 1000 docked structures by means of interface root mean square deviation (IRMSD) as a form of distance measurement. The third step eliminates steric overlaps with energy minimization. The output models thus obtained represent the models of the 10 most populated clusters and the centre, lowest energy weighted scores of 30 clusters along with cluster members are also displayed^44-46^. The selection of clusters for molecular dynamics simulation was performed based on both the no. of cluster members and weighted score of lowest energy.

### Binding affinity analysis

The binding affinity of the docked complexes were estimated with the binding free energy of the complex on the PROtein BinDIng enerGY prediction (PRODIGY) web server (https://bianca.science.uu.nl/prodigy/). It predicts the binding free energy of the biological complex. It incorporates predictive model solely based on intermolecular contacts along with non-interface surface properties. This server prediction methods have been compiled with Python scripts and Perl. The model predicts with accuracy on a huge heterogenous data showing a Pearson’s Correlation of 7.3, with p-value < 0.0001, between the predicted and experimental values and a Root Mean Square Error of 1.89 kcal mol^-1^. This server takes PDB input of docked complex, chain identifiers of the interacting molecules and temperature at which dissociation constant is measured The output results show the predicted binding free energy of complex (in kcal mol^-1^), dissociation constant (in M), the number as well as type of intermolecular contacts within 5.5 Å distance threshold and the percentages of charged along with polar amino-acids on the non-interacting surface^47,48^.

### Molecular Dynamics simulation of the vaccine construct

Molecular dynamics (MD) simulation of the vaccine construct was performed by Galaxy server^49^ which uses GROMACS (GROningen MAchine for Chemical Simulations) engine for running the simulation. MD simulation was done using TIP3P (Transferable Intermolecular potential with 3 points) water model and OPLS/AA (Optimized Potential for Liquid Simulation-All Atom) force field constraints. Solvation was done using Statistical Process Control (SPC) generic three-point model and charged ions were added for making the system neutral^50^. Energy minimization, NVT equilibration and NPT equilibration were performed for 100 ps (50,000 steps with 0.002 ps timestep). Production simulation of energy minimized structure was performed for 10 ns (5 times each of 2 ns using checkpoint files) using steepest descent algorithm, fast smooth particle-Mesh Ewald (SPME) electrostatics. Coulomb and Van der Waal’s cut-off were set at 1.0. Output trajectories from each run were combined by Visual Molecular Dynamics (VMD) software^51^. RMSD, RMSF, principal component analysis (PCA) plot and dynamical cross-relational matrix (DCCM) analysis were done using Bio3D^52^. GROMACS energy plots of temperature, pressure and density were also drawn by using Xmgrace graph plotting software^53^.

### Revisiting protein structure validation after MD simulation

Resulting output PDB structure file generated after molecular dynamics simulation of vaccine construct was modified for tertiary structure validation by MolProbity server^54,55^. Ramachandran plot analysis was performed again with RAMPAGE. Finally, ProSA – web and ERRAT were used to obtain the Z-score and quality factor to estimate errors in the structure.

### Codon optimization and in silico cloning of the vaccine

Codon optimization is a pre-requisite for effective cloning strategy of the final vaccine construct. The JCat tool (http://www.jcat.de/) was utilized for codon optimization and reverse translation to ensure expression of the construct in an expression vector as per the usage of the expression system. The tool takes protein sequence and the organism for vaccine expression, as input. The output includes sequence evaluation, Codon Adaptation Index (CAI), GC content of the sequence, adapted sequence. CAI score >0.8 and GC content in the range 30-70% assures efficient translation of the protein in the expression system^56,57^. This tool is amalgamated to the PRODORIC database which harbours all related data of different organisms. The SnapGene software was finally used to design the recombinant plasmid. Restriction sites of EcoRI and EcoRV were included at the 5’ and 3’ end of the vaccine construct respectively. This optimized the vaccine construct sequence comprising restriction sites, was cloned in pET-28a (+) vector using the software to assure expression.

### Immune Simulation

The assessment of the immunogenic profile of final vaccine construct was done with the C ImmSim web server (http://kraken.iac.rm.cnr.it/C-IMMSIM/index.php?page=1) for simulation of the immune response. This server incorporates prediction methods along with Miyazawa and Jernigan protein-protein potential measurements for the purpose of assessment of molecular binding in reference to immune complexes. A classical immunization experiment (entirely simulated for the model generated) reproduces development of immunological memory. At the end, the aspect of chronic exposure to the same immunogenic molecule leading to the emergence of one or more dominating clones of lymphocytes is even explored and the results manifest high affinity clones undergoing proliferation more than any other. The prediction methods employ algorithms which delineate biological complexes in the form of bit strings and Position Specific Scoring Matrix based methods for performance prediction. The server for simulation accepts, antigen sequence as input along with the matrices which define the binding motifs for the haplotype (four matrices for class I, two HLA-A and two HLA-B, as well as two matrices for class II) and other variables as well as parameters used to modulate the system^58^. The total time step of injection was set to 1400 (1-time step corresponds to 8 hour). The vaccine was injected at an interval of four weeks at following time steps 10, 94, 178, 262, 346, 430, 514, 598, 682, 766, 850, 934. After vaccination, a live replicating virus was injected at time step 1,100 with multiplication factor (0.2) and infectivity (0.6). As a control experiment, a live replicating virus was injected at the same time-step 1,100 without prior vaccination.

## Data availability

The datasets generated during and/or analysed during the current study are available from the corresponding author on reasonable request.

## Author contributions

SB and KM designed the protocols and the experiments. Multiple sequence alignments and binding analysis were done by BM. Tertiary structure prediction and docking experiments were done by KM. SB performed the physicochemical properties prediction, secondary structure prediction, tertiary structure refinement, codon optimization and cloning experiments. Epitope prediction, antigenicity prediction, allergenicity prediction and manuscript reviewing were done by GJG, DG and KM. SB performed the Molecular Dynamics simulation of the vaccine construct. Immune simulation was done by BM, SB and KM. Original manuscript has been written by KM. SB and BM have supervised the experiments. Final manuscript reviewing was done by all the authors.

## Additional information

