## Supplementary File for "Immuno-informatics approach for multi-epitope vaccine designing against SARS-CoV-2"

**Supplementary Table S1.** Prediction scores of selected CTL epitopes and affinities of selected HTL epitopes.

| Protein | CTL Epitopes | Combined Score | HTL Epitopes | Affinity (nM) |
|---|---|---|---|---|
| Nucleocapsid Phosphoprotein | GTDYKHWPQ | 1.21 | ASWFTALTQHGKEDL | 30.1 |
| | LLNKHIDAY | 1.39 | QIGYYRRATRRIRGG | 7.9 |
| | LSPRWYFYY | 2.34 | NNAAIVLQLPQGTTL | 10.6 |
| | | | HWPQIAQFAPSASAF | 17.2 |
| | | | QQTVTLLPAADLDDF | 7 |
| Membrane Glycoprotein | LLEQWNLVI | 0.77 | SYYKLGASQRVAGDS | 3.3 |
| | LVGLMWLSY | 1.40 | | |
| | YSRYRIGNY | 1.66 | | |
| Envelope Protein | VSLVKPSFY | 1.71 | | |
| ORF6 | NLDYIINLI | 0.79 | ILLIIMRTFKVSIWN | 5.8 |
| | LTENKYSQL | 0.95 | | |
| ORF7a | ITLATCELY | 1.42 | VKHVYQLRARSVSPK | 5.6 |
| | RQEEVQELY | 1.60 | | |
| ORF10 | QVDVVNFNL | 0.85 | | |

**Supplementary Table S2.** Discontinuous B-cell epitopes provided with prediction scores along with number of residues.

| Discontinuous Epitopes | Number of Residues | Score |
|---|---|---|
| VF (8-9), TVLLSSAY (11-18), E (32), NTQIHTLNDKIFSYTESLA (35-53) K (55), NGATFQVEVPGSQHIDSQKKAIERMKDTLRIAYLTEAKVEKLCVW (65-109) | 76 | 0.755 |
| AHGTPQN (19-25) | 7 | 0.719 |
| NLGPG (269-273), GYYRRATRRIR (298-308), PQGTTLGPGPGHWPQ (325-339), Q (342), PAADLDDF (363-370), YYKLGASQRVAGDSGPGPGILLIIMRTFKVSIWNGPGPGVKHVYQLRARSVSPKHHHHHH (377-436) | 100 | 0.702 |
| AKGTDYKHWPQ (128-138), AY (140-141), NK (144-145), L (167), Q (169) | 17 | 0.684 |
| PSASAFGPGPG (345-355) | 11 | 0.631 |
| RIG (194-196) | 3 | 0.615 |
| YSQLA (231-235) | 5 | 0.566 |
| MIK (1-3) | 3 | 0.531 |
| E (252), VQE (254-256), PGA (274-276), FT (279-280), TQHGK (283-287) | 14 | 0.518 |
| SLVK (203-206) | 4 | 0.505 |

**Supplementary Table S3.** Linear B-cell epitopes predicted by ElliPro with prediction scores along with number of residues.

| Linear B-cell Epitope | Number of Residues | Score |
|---|---|---|
| VFFTVLLSSAYAHGTPQN (8-25) | 18 | 0.763 |
| KLGASQRVAGDSGPGPGILLIIMRTFKVSIWNGPGPGVKHVYQLRARSVSPKHHHHHH (379-436) | 58 | 0.764 |
| NGATFQVEVPGSQHIDSQKKAIERMKDTLRIAYLTEAKVEKLCVWNN (65-111) | 47 | 0.762 |
| AKGTDYKHWPQAAY (128-141) | 14 | 0.725 |
| YYRRATRRIR (299-308) | 10 | 0.726 |
| NTQIHTLNDKIFSYTESL (35-52) | 18 | 0.704 |
| PQGTTLGPGPGHWPQ (325-339) | 15 | 0.67 |
| PSASAFGPGPG (345-355) | 11 | 0.631 |
| SQLA (232-235) | 4 | 0.584 |
| PAADLDDFG (363-371) | 9 | 0.55 |
| MIKLK (1-5) | 5 | 0.54 |
| TQHGK (283-287) | 5 | 0.529 |
| YRIG (193-196) | 4 | 0.519 |
| SLVK (203-206) | 4 | 0.505 |

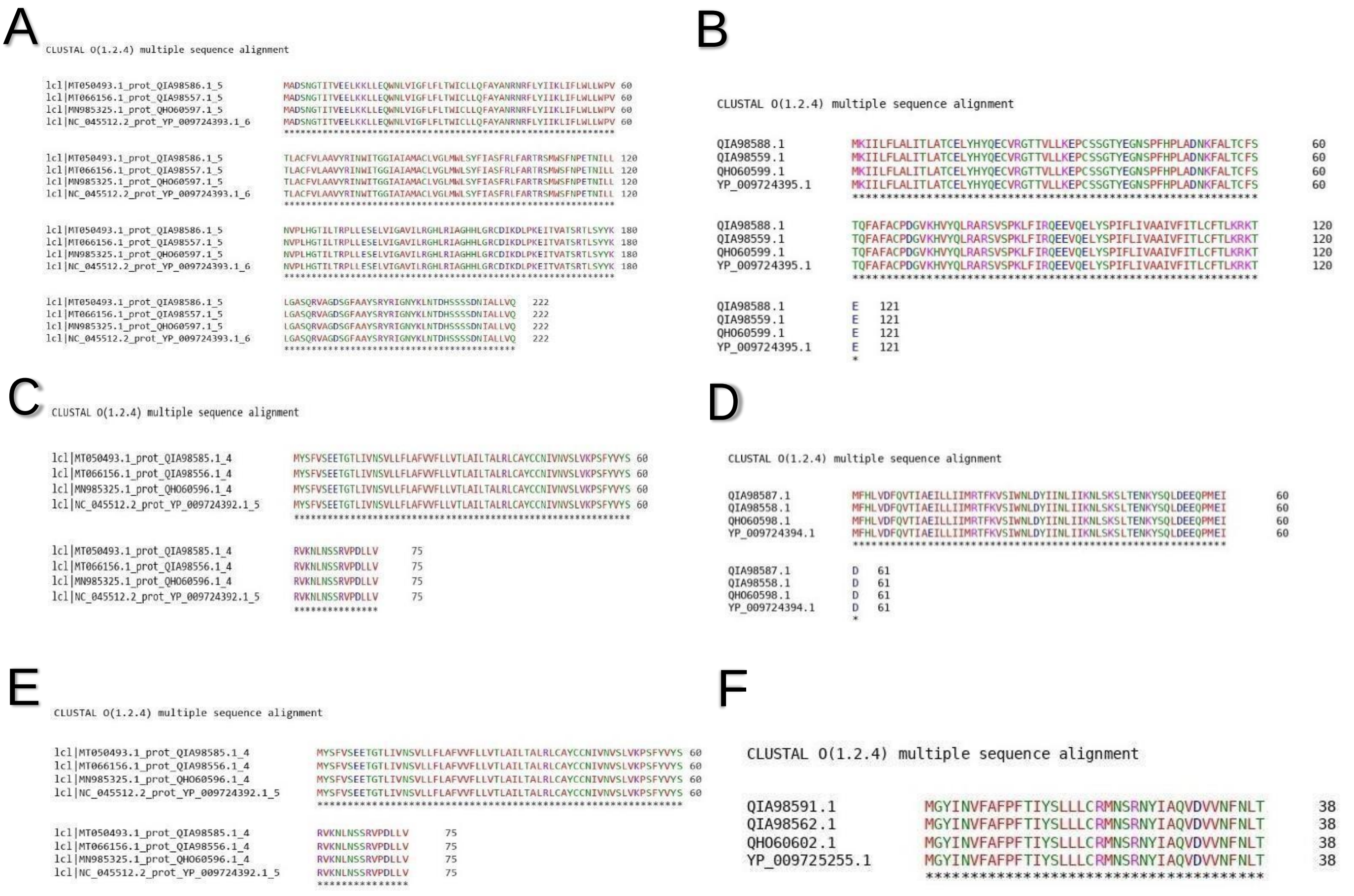

**Supplementary Figure S1. Multiple sequence alignment results of viral proteins**. **(A)** Nucleocapsid phosphoprotein **(B)** ORF7a **(C)** Envelope protein **(D)** ORF6 **(E)** Membrane glycoprotein **(F)** ORF 10.

Asterisk (*) represents conserved amino acid of each protein sequence in the four different viral strains.

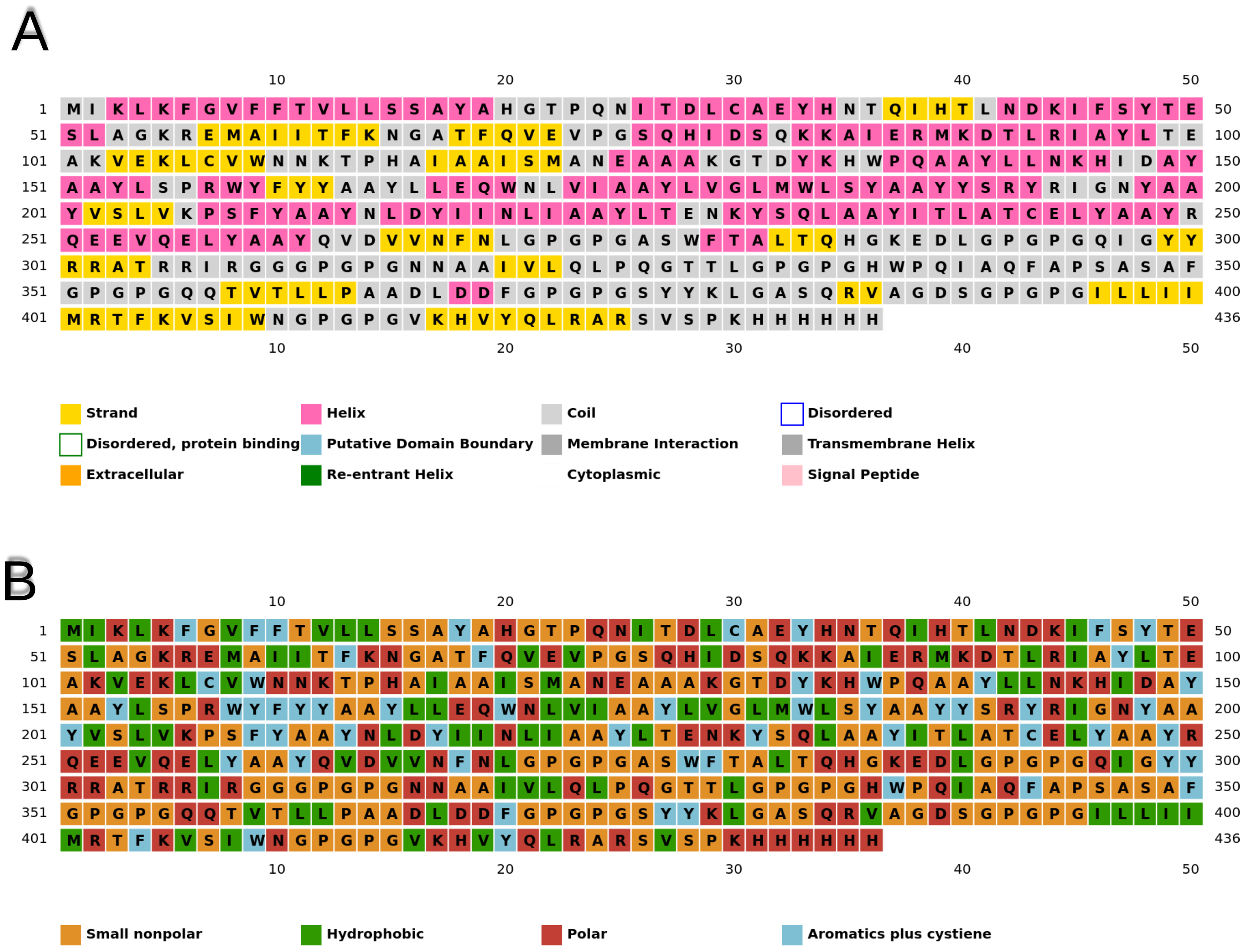

**Supplementary Figure S2. PSIPRED results. (A)** Results of secondary structure predicted by PSIPRED. **(B)** Chemical properties of each amino acid residue of the vaccine construct predicted by PSIPRED.

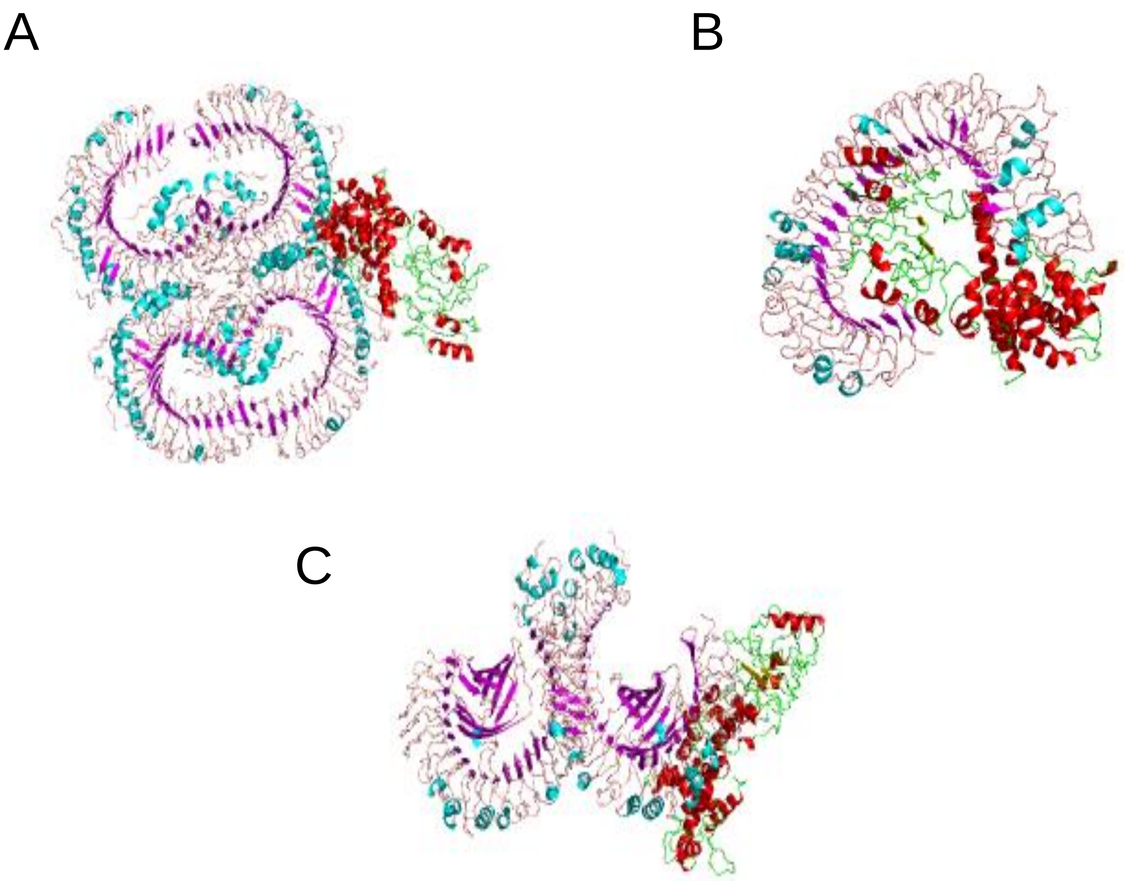

**Supplementary Figure S3. 3D visualization of docked complex in ClusPro. (A)** TLR2 - vaccine docked complex **(B)** TLR3 -vaccine docked complex and **(C)** TLR4 - vaccine docked complex. The beta-strands, alpha-helix, coils are indicated in the vaccine with brown, red and green colors respectively. The beta-strands, alpha-helix, coils are indicated in the receptor with pink, blue and pale brown colors respectively.

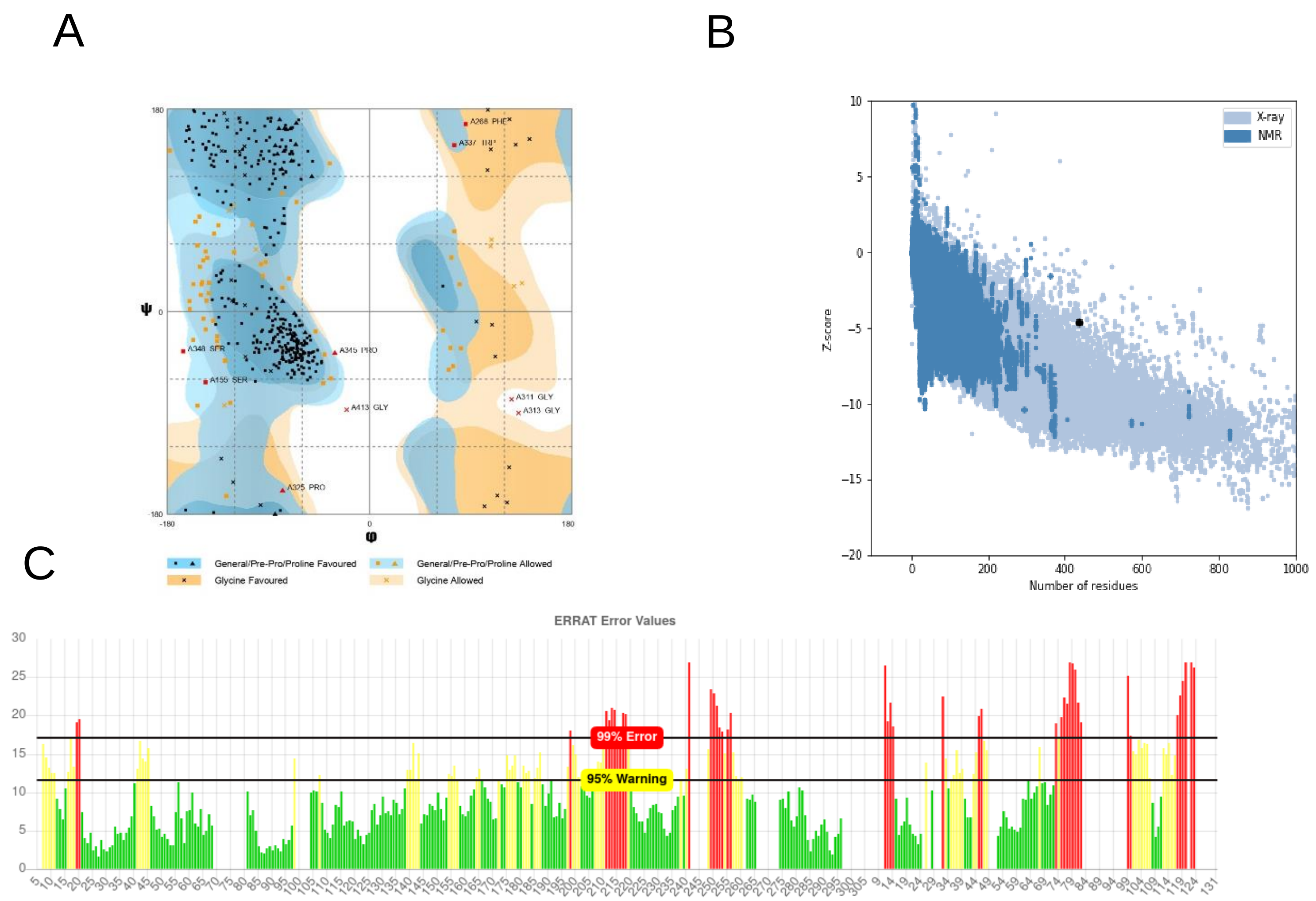

**Supplementary Figure S4. Protein tertiary structure validation after MD simulation.** (**A**) Ramachandran plot was analysed indicating 83.8%, 14.1% and 2.1% in favored, allowed and outlier regions respectively (**B**) Validation with ProSA-web providing a z score of -4.58. The black colored spot in the plot indicates z score. (**C**) Validation by ERRAT with quality factor 64.74.

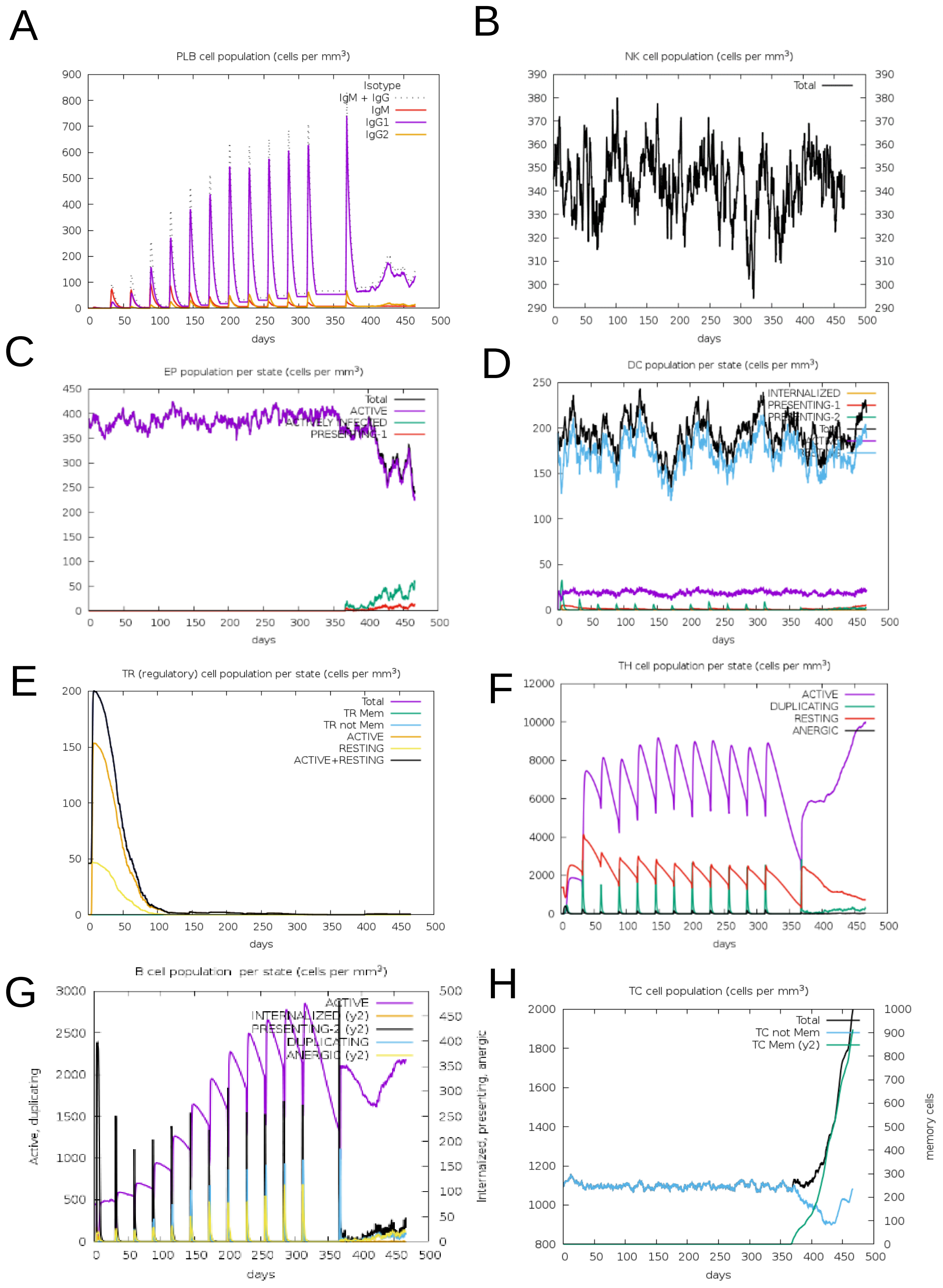

**Supplementary Figure S5. Immune simulation results of vaccine**. 12 doses of vaccine injections were given for almost 15 months and a live replicating virus was injected at around day 366. (**A**) evolution of plasma B cell population after each dose of injection. (**B**) consistency of NK cell population activity. (**C**) active epithelial cells are shown. (**D**) activation of dendritic cell population per state (**E**) regulatory T cells per state (**F**) resting state (shown in red) of helper T cell ($T_H$) represents cells not presented to the antigen and active state (shown in violet) represents activated $T_H$ cells in response to the antigen (**G**) activation of B cell population per state and (**H**) evolution of cytotoxic T cell population clearly shows the activity of memory cells developed after virus injection due to prior vaccination.
